# Optimizing phage-antibiotic combinations: impact of administration order against daptomycin non-susceptible (DNS) MRSA clinical isolates

**DOI:** 10.1101/2025.05.19.654851

**Authors:** Callan R. Bleick, Sean Van Helden, Andrew D. Berti, Rita Richa, Susan M. Lehman, Arnold S. Bayer, Michael J. Rybak

**Affiliations:** Anti-Infective Research Laboratory, Department of Pharmacy Practice, Eugene Applebaum College of Pharmacy and Health Sciences, Wayne State University, Detroit, MI, USA; Center for Biologics Evaluation and Research, US Food and Drug Administration, Silver Spring, MD, USA; The Dept of Medicine, David Geffen School of Medicine at UCLA, Los Angeles, CA, USA; The Lundquist Institution for Biomedical Innovation at Harbor-UCLA, Torrance, CA, USA; Department of Pharmacy Services, Detroit Receiving Hospital, Detroit Medical Center, Detroit, MI, USA; Department of Medicine, Division of Infectious Diseases, Wayne State University, Detroit, MI, USA

**Keywords:** Bacteriophage therapy, antibiotic resistance, phage-antibiotic interactions, PACs, combination therapy, order effects, administration timing, DNS, MRSA

## Abstract

**Background:** The rise of bacterial resistance has driven the exploration of novel therapies, such as bacteriophage-antibiotic cocktails (PACs), which have shown in vitro promise against resistant pathogens, including daptomycin non-susceptible (DNS-MRSA) strains. While daptomycin has been a cornerstone for treating MRSA bacteremia and vancomycin-refractory infective endocarditis, the emergence of DNS-MRSA presents a significant challenge due to high morbidity, mortality, and rapid intrinsic resistance development.

**Methods:** Phages Intesti13 and Sb-1, were selected for their unique host ranges and activity against sixteen DNS-MRSA strains. Synergy with antibiotics was assessed via growth suppression curves and 24-hour time-kill assays (TKAs) across administration sequences and MIC increments. Selected regimens were further assessed in an *ex-vivo* simulated endocardial vegetation (SEV) models, with pharmacokinetic analyses confirming target antibiotic concentrations.

**Results:** In the *ex-vivo* SEV model, simultaneous PAC administration using daptomycin±phage, showed superior bactericidal activity over sequential treatments in isolate C6 (p<0.01). Similarly, in the same model, C2 reached detection limits within 48h and remained suppressed for 120h (p<0.0037). Sequential outcomes varied by phage-antibiotic order and antibiotic choice. Simultaneous and phage-first regimens outperformed antibiotic-first, especially in 24h TKAs, but showed variability at lower MICs and between *in-vitro* and *ex-vivo* settings.

**Conclusion:** This study highlights PAC’s potential for DNS-MRSA treatment, emphasizing the importance of administration timing. The observed differences across clinical strains emphasize the need for strain-specific evaluations and a deeper understanding of phage-antibiotic interactions to optimize therapy. Future research must focus on expanding phage diversity, refining protocols, and clinically validating sequential strategies to enhance PAC efficacy.

## INTRODUCTION

Patients with persistent methicillin-resistant *Staphylococcus aureus* (MRSA) infections, which are associated with high mortality rates, often require prolonged hospitalizations, intensive care services, extended use of costly alternative antimicrobial therapies, and increased treatment for disease-related complications. (1, 2), (3), (4) Addressing these challenges remain a critical concern in both clinical and public health settings Daptomycin has been a cornerstone treatment for the management of MRSA bacteremia and infective endocarditis infections refractory to vancomycin. (5, 6) However, the emergence of daptomycin non-susceptible (DNS) MRSA presents a significant treatment challenge, as these infections have been associated with high morbidity and mortality due to difficulties in achieving effective eradication along with acquired resistance to antibiotics. (7–9)

In the United States, DNS-MRSA is classified as a serious threat, with an estimated 323,700 cases annually, resulting in 10,600 deaths and $1.7 billion in healthcare costs. (4) This concern is especially alarming in immunocompromised patients and those with comorbidities, where treatment failure can result in even more severe consequences. (2) Recent studies have identified reduced susceptibility to daptomycin in a subset of patients with MRSA endocarditis (6), while others highlight the role of DNS-MRSA variants in driving resistance. (10, 11) The loss of vancomycin and daptomycin as reliable first-line therapeutic agents further narrows the already limited treatment options available for these infections.

Tackling these issues requires a multifaceted approach that focuses on innovative research aimed at overcoming the challenges posed by antimicrobial resistance (AMR). Given the urgent need to explore alternative strategies, bacteriophage (phage) therapy has emerged as a promising therapeutic approach targeting these difficult-to-treat infections. (12–15) This novel approach may play a crucial role in conquering the growing challenges arising from AMR.

Obligately lytic phages are prevalent in nature and are highly-selective predators of bacteria which have demonstrated promising results when used in combination with antibiotics. (13, 14, 16–19) The benefits of utilizing phage in combination with antibiotics include synergistic bacterial killing, improved antimicrobial penetration, restoration of antibiotic activity in resistant strains, and eradication of bacterial biofilm. (16, 18, 20) While controlled clinical data are lacking, many in vitro experiments have shown that combinations of phages and antibiotics can be more effective than antibiotics alone. (21) Prior studies from our group have demonstrated potent, synergistic bactericidal activity against DNS-MRSA clinical isolates when a cocktail of phages was added to antibiotic therapy under humanized antibiotic exposures. (17, 22) Other studies have also shown that combining phages with antibiotics results in synergistic effects, characterized by bigger plaque sizes and improved lytic activity. (15, 23, 24) However, there is often substantial variability among bacterial strains in terms of which combinations of phages and antibiotics have additive or synergistic effects. In our own studies, we have used a combination of planktonic time-kill assays, biofilm time-kill assays, and more complex two-compartment models for longer-term experiments that mimic human pharmacokinetic/pharmacodynamic parameters. Collectively, our data support a preliminary hypothesis that a triple combination of i) daptomycin or vancomycin, ii) a β-lactam such as ceftaroline or cefazolin, and iii) a cocktail of staphylococcal myophages can overcome much of this strain variability. (17, 21, 25, 26)

Most of our experiments have involved simultaneous initiation of phage and antibiotic treatment, which might not be representative of many clinical scenarios as patients using phages in clinical trials or under expanded access are likely to already be receiving antibiotics.

Therefore, we want to understand the impact that the sequence of phage and antibiotic administration might have. Some studies have explored this question, but much remains unclear. (27–29) Improving our understanding of the temporal dynamics between antibiotics and phage is crucial in maximizing their therapeutic potential. In this study, we aim to investigate the effect on bacterial eradication when administering phages simultaneous to antibiotics versus sequentially alongside antibiotics.

## RESULTS

### Continuous Growth Suppression Screening over 24h

We used continuous growth suppression screening identified the strains that were least susceptible to the daptomycin-ceftaroline (DAP+CPT) combination. The culture turbidity [OD_600_] of the 16 DNS-MRSA isolates in the presence of DAP+CPT at 0.5x MIC was monitored over 24h. All isolates shown in Figure 1A as well as JKD6005, C27, and C39 shown in Figure 1B, were poorly suppressed, with OD_600_ values exceeding the growth suppression threshold before 24h. The combination of DAP+CPT at 0.5x MIC was the most effective against strains J03, C21, C43, C18, and C51 represented in Figure 1B, maintaining OD_600_ values below the growth suppression threshold (< 0.26) for up to 23h, reflecting significant suppression compared to all other isolates at 24h (p < 0.007, 0.012).

**Figure 1.**
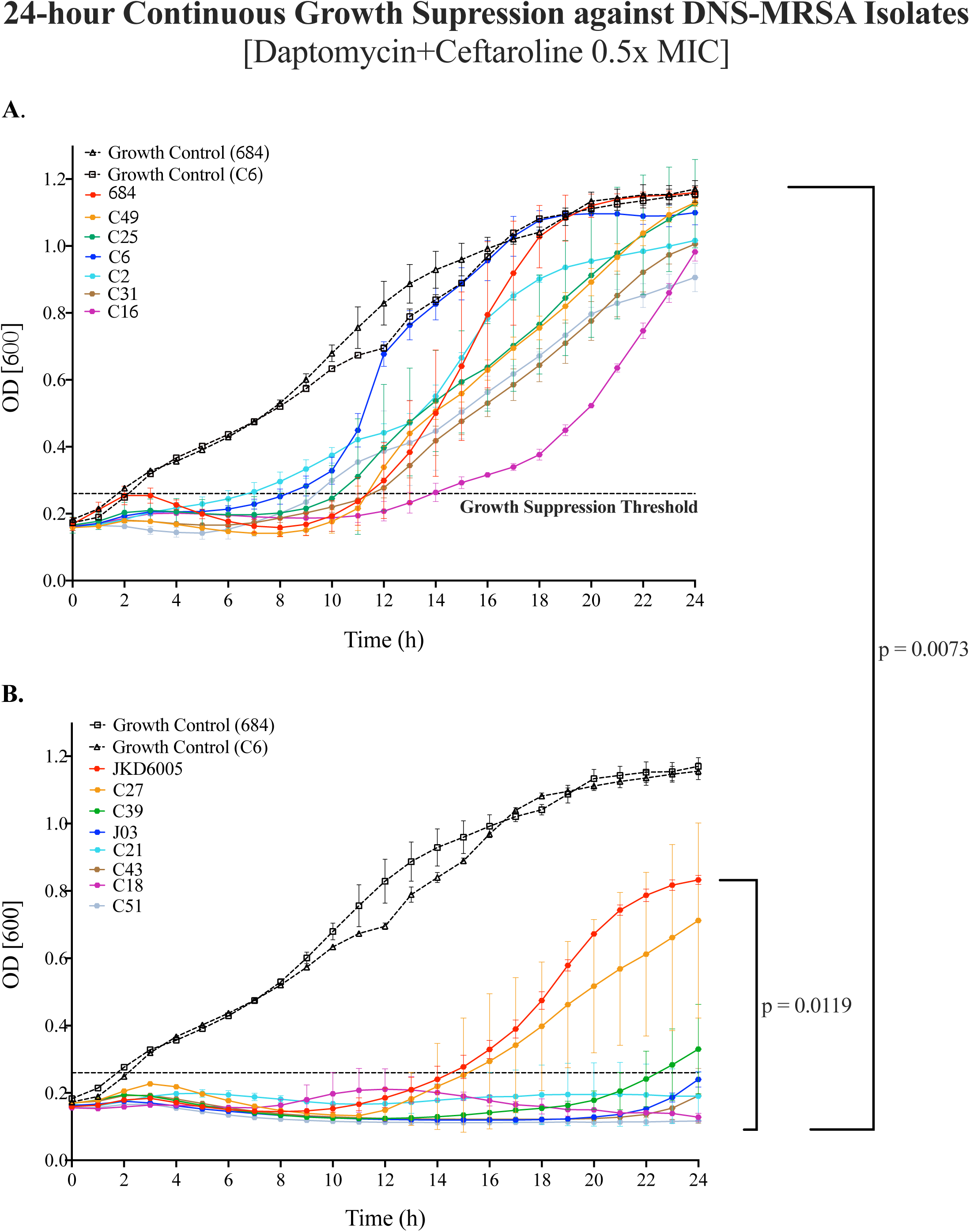
**(A, B)**: 24h continuous growth suppression of DNS-MRSA isolates (n=16) against daptomycin (DAP) and ceftaroline (CPT) at 0.5x MIC. Bacterial growth was monitored via OD [600] over 24h following treatment with sub-inhibitory daptomycin and ceftaroline (0.5x MIC). Strains are ranked in the legend by OD [600] at 24h (highest to lowest). A growth suppression threshold of 0.266 (95% CI lower bound) was used to identify significantly suppressed isolates. Statistical analysis included two-way repeated measures ANOVA with Geisser-Greenhouse correction and Tukey9s HSD post hoc test (p<0.005) for multiple comparisons. Significant interactions at individual time points were further examined (S1, S2). *See Supplementary Tables S1 and S2 for all Tukeys9s pairwise comparisons at T0 and T24*.

These findings highlight the variability in growth dynamics among bacterial isolates exposed to subinhibitory concentrations of DAP+CPT over 24h. All pairwise comparisons are provided in are provided in Supplementary Tables S1 and S2.

### 24h Time-kill Analyses: Antibiotics Alone

Based on growth suppression data, five poorly-suppressed isolates, defined by the mean upper 95% confidence interval across all strains OD_600_ ≥ 0.42, were selected to progress to time-kill assays (TKAs) for further analysis. These isolates included C31, C49 (USA300/ST8 lineage), C2, C6 (USA100/ST5 lineage) and 684 (USA200/ST30 lineage). The results in Figure 2 A-E demonstrate the efficacy of antibiotic alone and in combination across varying antibiotic concentrations (0.5x, 1x, 2x MIC).

**Figure 2.**
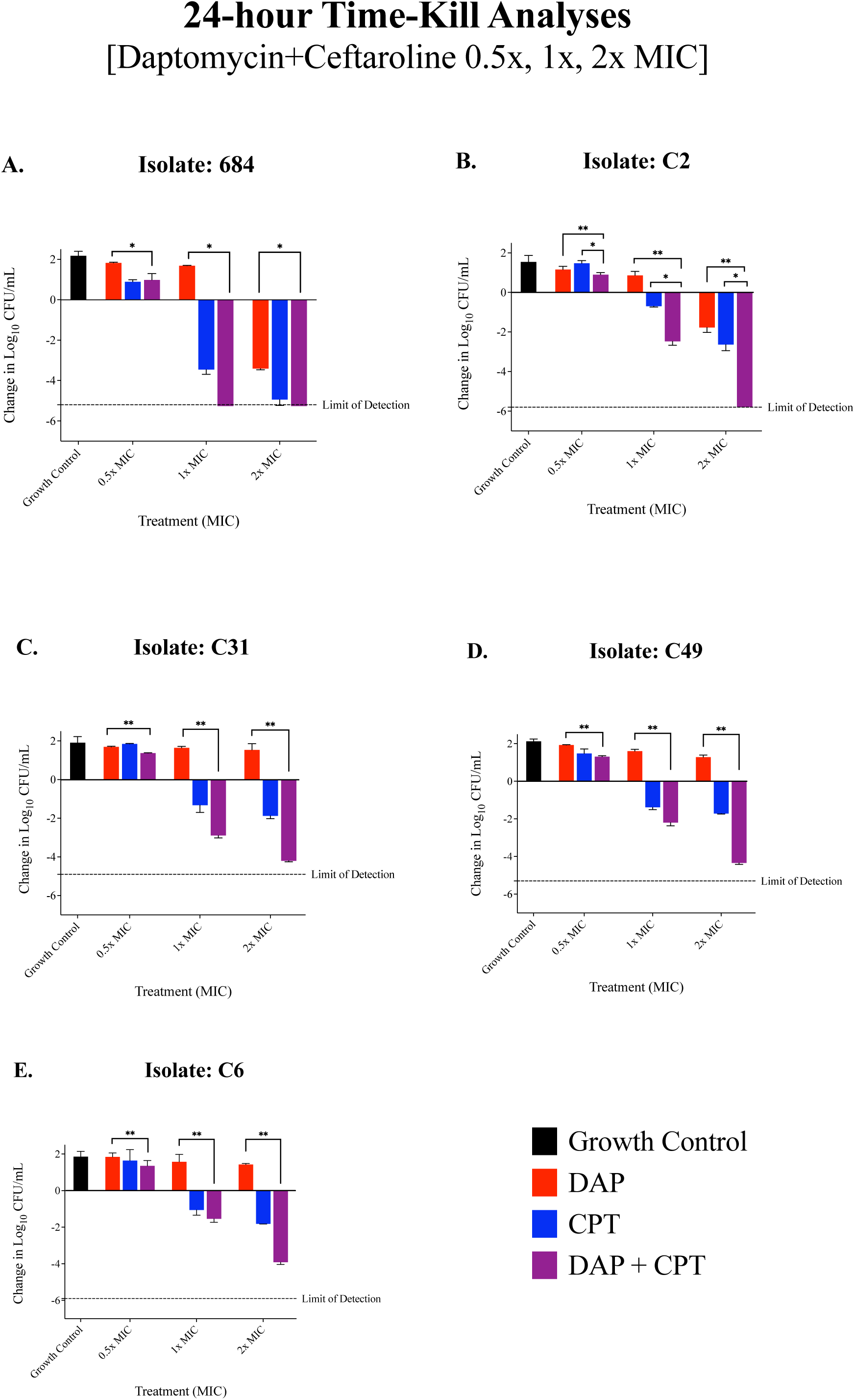
**(A-E):** 24-hour time-kill analyses conducted in a planktonic state against five S. aureus strains treated with daptomycin+/-ceftaroline at varying concentrations of the MIC to identify S. aureus strains least susceptible to the combination of DAP+CPT. The change in bacterial counts (log¡ CFU/mL) was assessed across different treatment groups: growth control, antibiotics (DAP+CPT) at 0.5x MIC, 1x MIC, and 2x MIC. Overall, isolates C31, C49, and C6 showed moderate bacterial reductions with DAP + CPT at 2x MIC, but these did not reach the limit of detection (−4.20 ± 0.05, −4.34 ± 0.07, and −3.91 ± 0.11 log¡ CFU/mL, respectively). ANOVA and post-hoc tests revealed dose-dependent responses across all tested concentrations. Across all isolates, Tukey’s post-hoc test consistently indicated significantly lower bacterial reductions at 0.5x MIC compared to 1x and 2x MIC, highlighting the enhanced efficacy of higher antibiotic concentrations in this combination therapy. Error bars represent standard deviation from the mean, and statistical significance was determined using ANOVA followed by post-hoc Tukey’s test, with p < 0.05 considered significant. The limit of detection is indicated by the dashed line (>2 log10 CFU/mL reduction from baseline). P-values are represented as follows: *p < 0.05, **p < 0.01, ***p < 0.001, and ****p < 0.0001. Non-significant differences are denoted as ’ns’ (p g 0.05). Statistical significance was determined using ANOVA followed by post-hoc Tukey’s test. *All Tukey9s pairwise comparisons, can be found in Supplementary Figure S3*.

At 0.5 x MIC, DAP±CPT had minimal to no inhibitory activity relative to the growth control, which was expected. Isolates 684 and C2 were reduced to the detection limit by 2x MIC and also by 1x MIC for 684.

### Effects of Sequential Phage and Antibiotic Administration

TKAs were conducted on isolates C6, C49, and C31. These isolates were selected due to poor susceptibility against daptomycin (DAP) and ceftaroline (CPT). Additionally, isolate C2, which is known to be susceptible to DAP+CPT without the addition of phage, was included in partial TKAs (using daptomycin alone) as a control. This allowed us to assess whether sequential dosing influences daptomycin activity under different conditions and compare its effects between susceptible and less susceptible isolates. Figure 3 outlines the various dosing regimens that were used to evaluate sequential administration. These evaluations were performed at three antibiotic concentrations: 0.5x MIC, 1x MIC, and 2x MIC. Changes in bacterial counts (log₁₀ CFU/mL) for each treatment regimen are detailed in Figures 4-7 and described below, by isolate.

**Figure 3.**
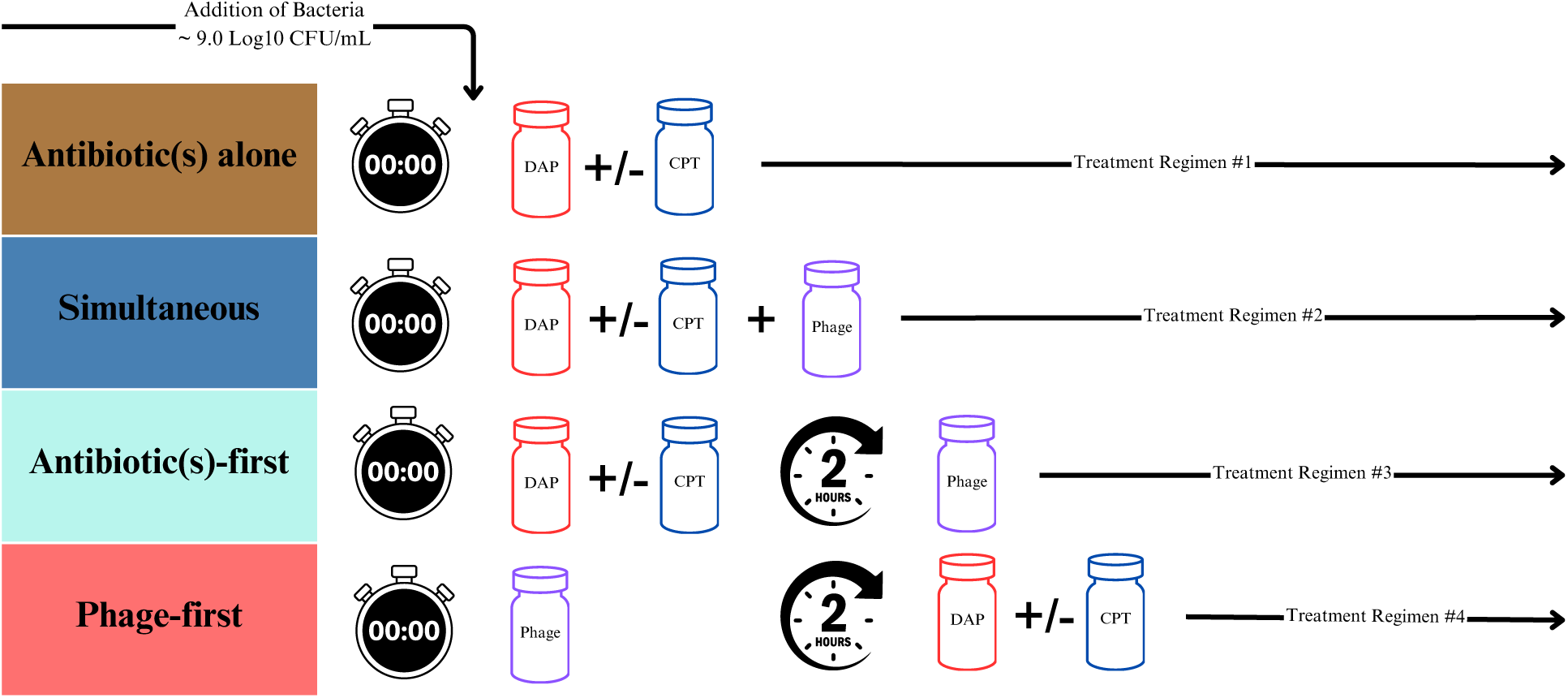
Schematic of dosing regimens across experimental settings The schematic above illustrates the different administration orders evaluated between two experimental settings: 24-hour time-kill assays and *ex-vivo* SEV models. Four distinct treatment regimens were evaluated, each allowing a 2-hour window from the initiation of the first dose: 1. “Antibiotics alone” - antibiotics were administered without any phage treatment. 2. “Simultaneous” - dosing of antibiotics and phage (blue): Both antibiotics and phage were administered at the same time. 3. “Antibiotic first” - dosing 2 hours before phage administration (aqua): Antibiotics were administered 2 hours prior to phage treatment. 4. “Phage first” - antibiotic dosing 2 hours after phage administration (red): Phage treatment was initiated 2 hours before the administration of antibiotics.

**Figure 4.**
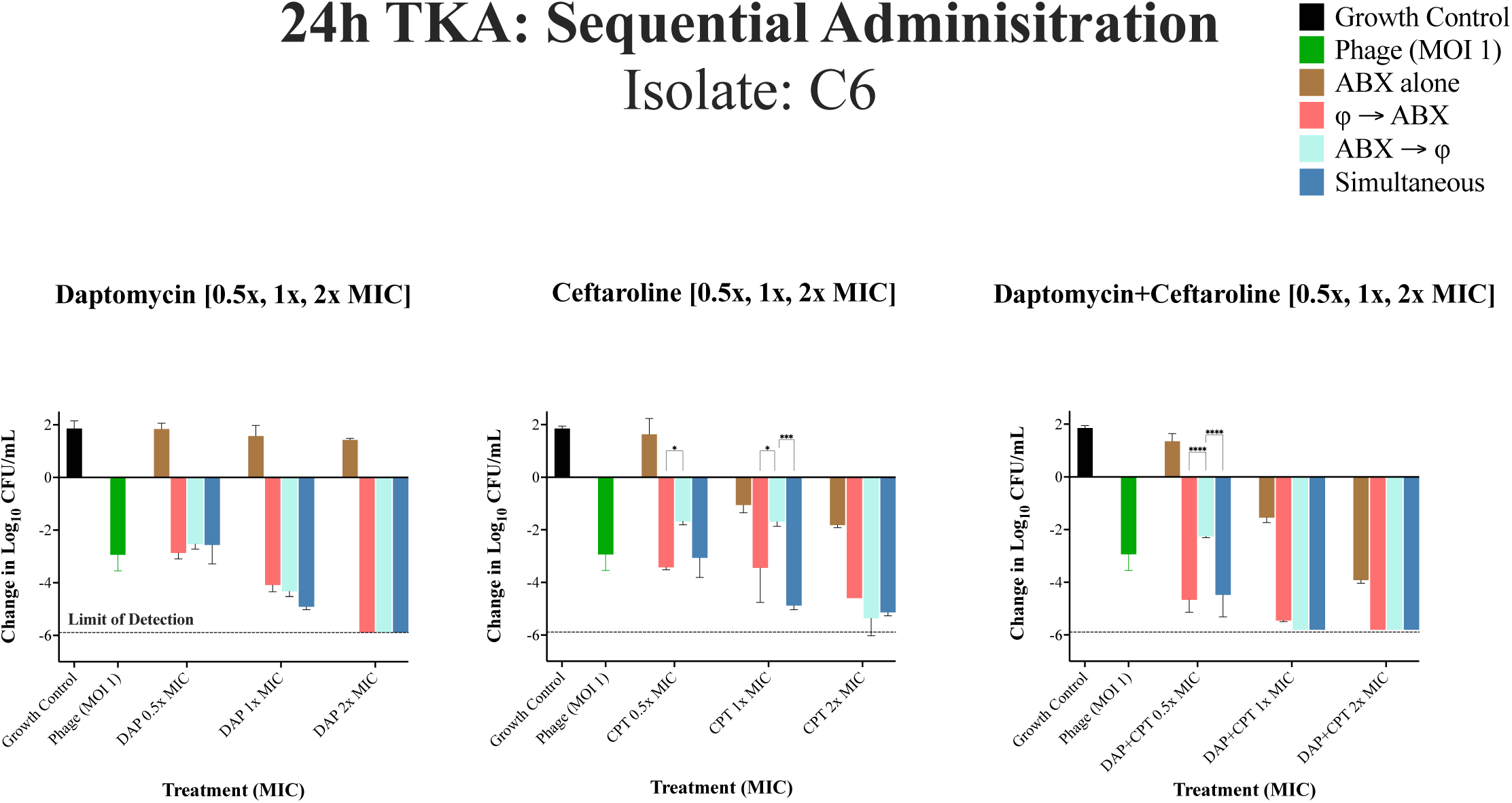
24h TKA for DNS-MRSA isolate C6 (USA100/ST5) against different administration regimens at 0.5x, 1x, and 2x MIC. Under Different Administration Regimens at 0.5x, 1x, and 2x MIC. Bacterial reductions were evaluated using a two-way repeated measures ANOVA to assess the effects of administration sequence (phage-first, antibiotic-first, simultaneous) and MIC levels (0.5x, 1x, and 2x MIC). Data were grouped by administration sequence (columns) and MIC levels (rows) to examine main and interaction effects. Pairwise comparisons were performed using Tukey9s HSD post hoc test. The dashed line represents the limit of detection. P-values are represented as follows: *p < 0.05, **p < 0.01, ***p < 0.001, and ****p < 0.0001. Non-significant differences are denoted as ’ns’ (p g 0.05). Statistical significance was determined using ANOVA followed by post-hoc Tukey’s test. Error bars show the standard deviation from the mean of biological replicates (conducted in duplicate).

### 24-hour Time Kill Assay: Sequential Phage and Antibiotic Administration against Isolate C6

In Figure 4 against C6, the phage cocktail alone had significant activity compared to the growth control (−2.94 log₁₀ CFU/mL ± 0.28, p < 0.0001). Administration of antibiotic alone demonstrated minimal to no activity compared to growth control, except at 2x MIC with DAP+CPT, which achieved a significant reduction (−5.81 log₁₀ CFU/mL ±0.11) compared to CPT (−1.82 log₁₀ CFU/mL ±0.01) or DAP alone (−1.4 log₁₀ CFU/mL ±0.05) at 2× the MIC (p < 0.001). The order of administration had no significant impact on bacterial eradication with DAP alone with phage, but increasing DAP concentration was associated with increased eradication across 0.5×, 1×, and 2× MIC levels (p < 0.001). For CPT-containing regimens (without DAP), phage-first or simultaneous administration significantly reduced CFU/mL at 0.5× and 1× the MIC compared to antibiotic-first administration (p < 0.001). Similarly, for DAP+CPT, these administration sequences were most effective at 0.5× the MIC (p < 0.001). However, the effect of sequential dosing diminished at higher concentrations, as reductions at 2× the MIC for CPT alone and 1× and 2× MIC for DAP+CPT approached or met the limit of detection, irrespective of the sequence of administration.

### 24-hour Time Kill Assay: Sequential Phage and Antibiotic Administration against Isolate C49

In Figure 5 against C49, the phage cocktail alone had minimal activity compared to growth control. The impact of sequential dosing was significant for both phage-first and simultaneous phage-antibiotic administration at 0.5x and 1x MIC, resulting in greater bacterial reductions compared to administering antibiotic first (p < 0.0001), with the exception of DAP+CPT at 1x and 2x MIC, where both antibiotic first or simultaneous dosing achieved the highest bacterial reductions (−1.56 log_10_ CFU/mL ± 0.06, p < 0.0001). There were no significant differences in bacterial reductions for DAP+CPT at 2x MIC. Using DAP alone at 2x MIC, simultaneous or administering antibiotics first prior to phage had the highest bacterial reductions compared to administering phage first (−1.07 log_10_ CFU/mL ± 0.12, p= 0.004).).

**Figure 5.**
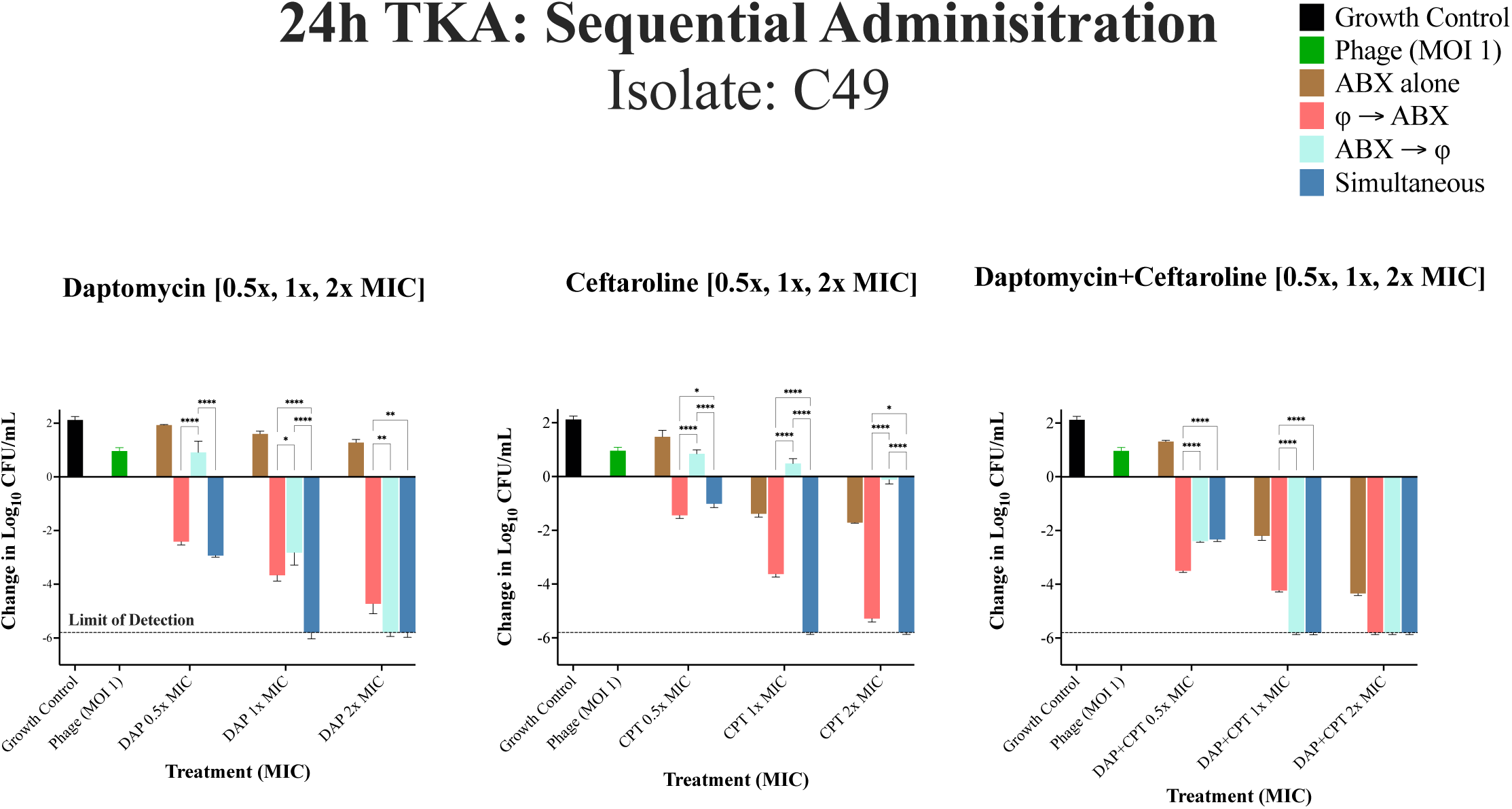
24h TKA for DNS-MRSA isolate C49 (USA300/ST8) against different administration regimens at 0.5x, 1x, and 2x MIC. Bacterial reductions were evaluated using a two-way repeated measures ANOVA to assess the effects of administration sequence (phage-first, antibiotic-first, simultaneous) and MIC levels (0.5x, 1x, and 2x MIC). Data were grouped by administration sequence (columns) and MIC levels (rows) to examine main and interaction effects. Pairwise comparisons were performed using Tukey9s HSD post hoc test. The dashed line represents the limit of detection. P-values are represented as follows: *p < 0.05, **p < 0.01, ***p < 0.001, and ****p < 0.0001. Non-significant differences are denoted as ’ns’ (p g 0.05). Statistical significance was determined using ANOVA followed by post-hoc Tukey’s test. Error bars show the standard deviation from the mean of biological replicates (conducted in duplicate).

### 24-hour Time Kill Assay: Sequential Phage and Antibiotic Administration against Isolate C31

In Figure 6 against C31, the phage cocktail alone had no meaningful reduction in log_10_ CFU/mL compared to growth control. Administration of antibiotics alone also displayed limited activity with DAP and CPT monotherapy. However, at 1x and 2x MIC of DAP+CPT, significant bacterial reductions were observed in the antibiotic alone regimen, outperforming phage-first administration (−1.57±0.12 vs. −2.21± 0.51 log₁₀ CFU/mL; p < 0.0001). In general, the order of administration had minimal to no impact in DAP alone regimens at 0.5x, 1x and 2x MIC while for CPT-only regimens, simultaneous or antibiotic first had a significantly greater reduction than phage first at 2x MIC. Antibiotic-first or simultaneous administration, followed by antibiotic alone regimens were significantly improved compared to phage-first administration for DAP+CPT at 2x the MIC (−2.81 log₁₀ CFU/mL ±0 .12, p < 0.0001).

**Figure 6.**
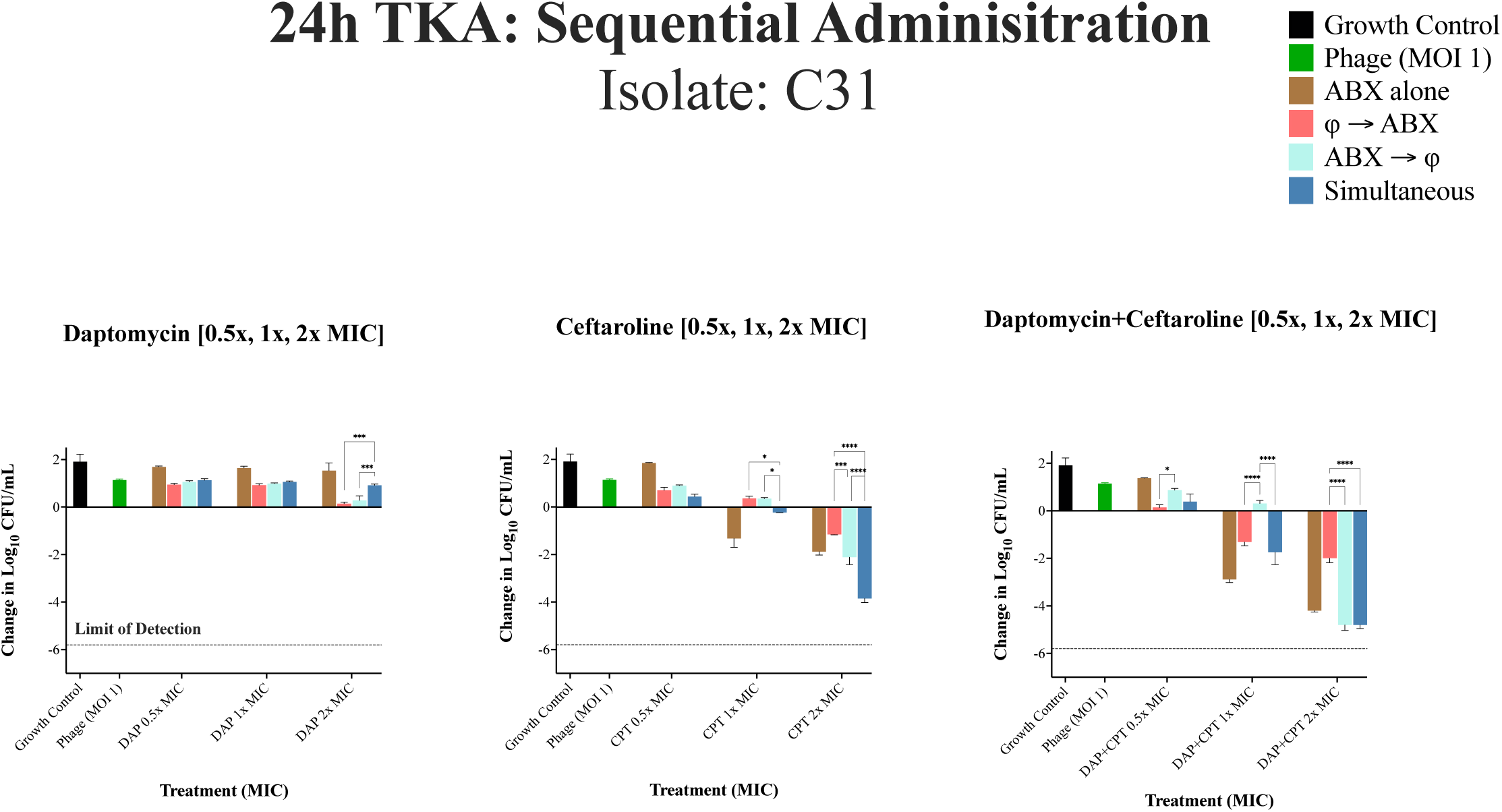
24h TKA for DNS-MRSA isolate C31 (USA300/ST8) against different administration regimens at 0.5x, 1x, and 2x MIC. Bacterial reductions were evaluated using a two-way repeated measures ANOVA to assess the effects of administration sequence (phage-first, antibiotic-first, simultaneous) and MIC levels (0.5x, 1x, and 2x MIC). Data were grouped by administration sequence (columns) and MIC levels (rows) to examine main and interaction effects. Pairwise comparisons were performed using Tukey9s HSD post hoc test. The dashed line represents the limit of detection. P-values are represented as follows: *p < 0.05, **p < 0.01, ***p < 0.001, and ****p < 0.0001. Non-significant differences are denoted as ’ns’ (p g 0.05). Statistical significance was determined using ANOVA followed by post-hoc Tukey’s test. Error bars show the standard deviation from the mean of biological replicates (conducted in duplicate).

**Figure 7.**
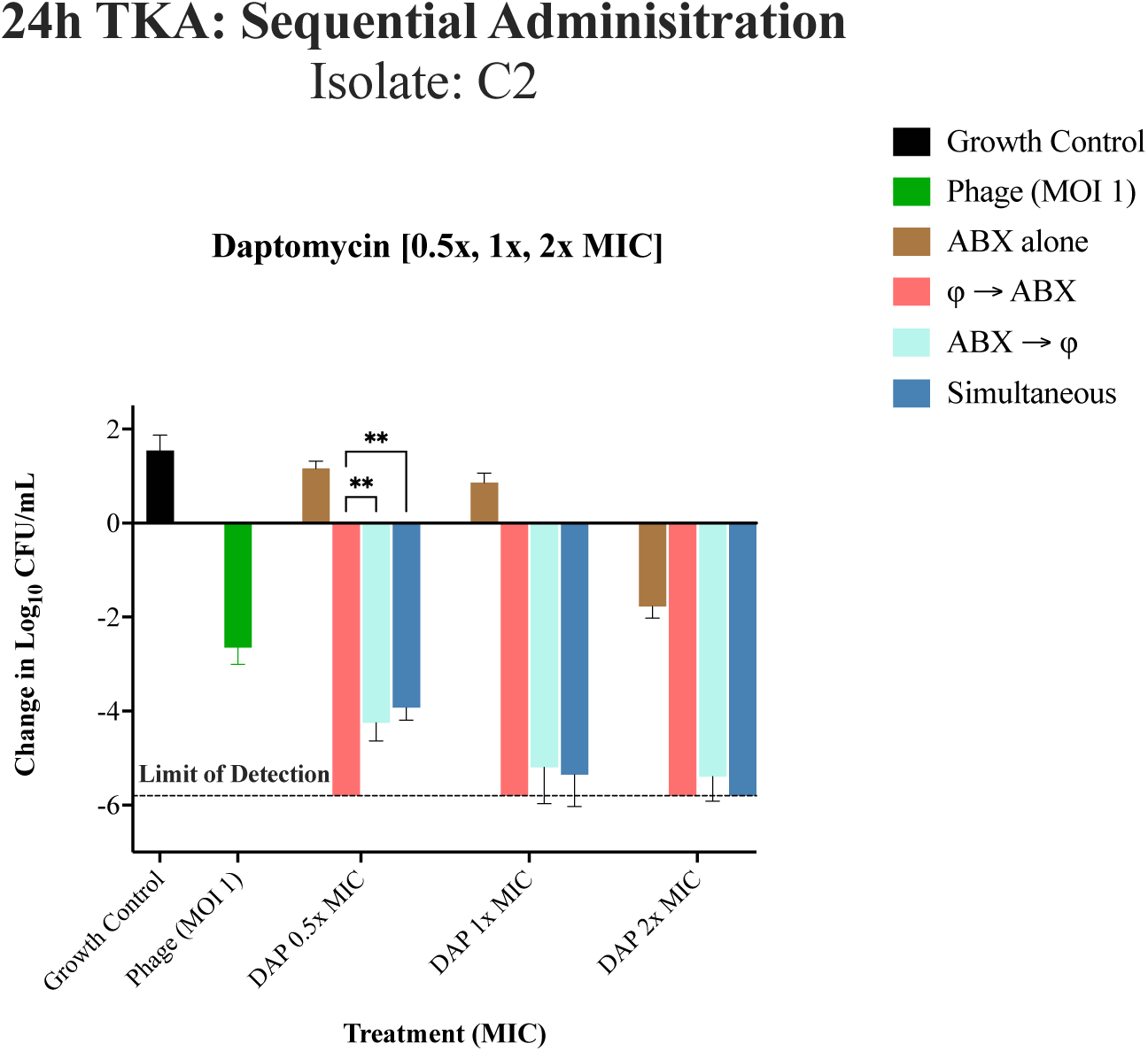
24h TKA for DNS-MRSA isolate C2 (USA100/ST5) against different administration regimens at 0.5x, 1x, and 2x MIC using DAP only. Bacterial reductions were evaluated using a two-way repeated measures ANOVA to assess the effects of administration sequence (phage-first, antibiotic-first, simultaneous) and MIC levels (0.5x, 1x, and 2x MIC). Data were grouped by administration sequence (columns) and MIC levels (rows) to examine main and interaction effects. Pairwise comparisons were performed using Tukey9s HSD post hoc test. The dashed line represents the limit of detection. P-values are represented as follows: *p < 0.05, **p < 0.01, ***p < 0.001, and ****p < 0.0001. Non-significant differences are denoted as ’ns’ (p g 0.05). Statistical significance was determined using ANOVA followed by post-hoc Tukey’s test. Error bars show the standard deviation from the mean of biological replicates (conducted in duplicate).

### 24-hour Time Kill Assay: Sequential Phage and Antibiotic Administration against Isolate C2

In Figure 7 against C2, the phage cocktail was significantly active compared to growth control and antibiotics alone at 0.5x the MIC (−2.5 ± 0.35 log₁₀ CFU/mL (p < 0.0001). With respect to order of administration, at 0.5x the MIC, phage administration first was significantly better than antibiotic first (−1.55 ±0.38 log₁₀ CFU/mL, p = 0.0062) or simultaneous administration (−1.87±0.26 log₁₀ CFU/mL, p = 0.0015). Since all of the treatment regimens using 1x or 2x MIC reduced bacterial populations to or below the limit of detection, no significant differences among them were detectable. This made it challenging to differentiate the effects of the administration sequences, except in the case of DAP alone.

### Ex-vivo Simulated Endocardial Vegetation (SEV) Model

We investigated four administration sequences using humanized doses of daptomycin (DAP) and/or ceftaroline (CPT) with a phage cocktail against DNS-MRSA isolates C6 and C2 at a high inoculum. C2 was selected for its strong phage susceptibility, while C6 exhibited poor response to the phage cocktail alone. A total of 40 unique *ex vivo* PK/PD SEV experiments were conducted in duplicate, including four treatment regimens, a growth control, and a phage-only group (Figures 8, 9). Humanized doses of DAP (10 mg/kg/day) were used with a total maximum drug concentration (C_max_) reaching 138.2 ± 0.44 μg/mL (target total C_max_, 141.1 μg/mL) and half-life (*t_½_*) of 7.7 ± 0.17 h (target *t*_½_, 8 h) and an area under the curve (AUC) from 0 to 24 h of 1,714.88 ± 5.2 μg · h/mL (Table 2). The PK parameters for CPT in the model consisted of a C_max_ of 20.15 ± 0.54 μg/mL (target total C_max_, 21.3 μg/mL), a *t_½_* of 2.54 ± 0.08 h (target *t_½_*, 2.66 h), and an AUC from 0 to 24 h (AUC_0–24_) of 67.40 ± 8.61 μg · h/mL.

**Figure 8.**
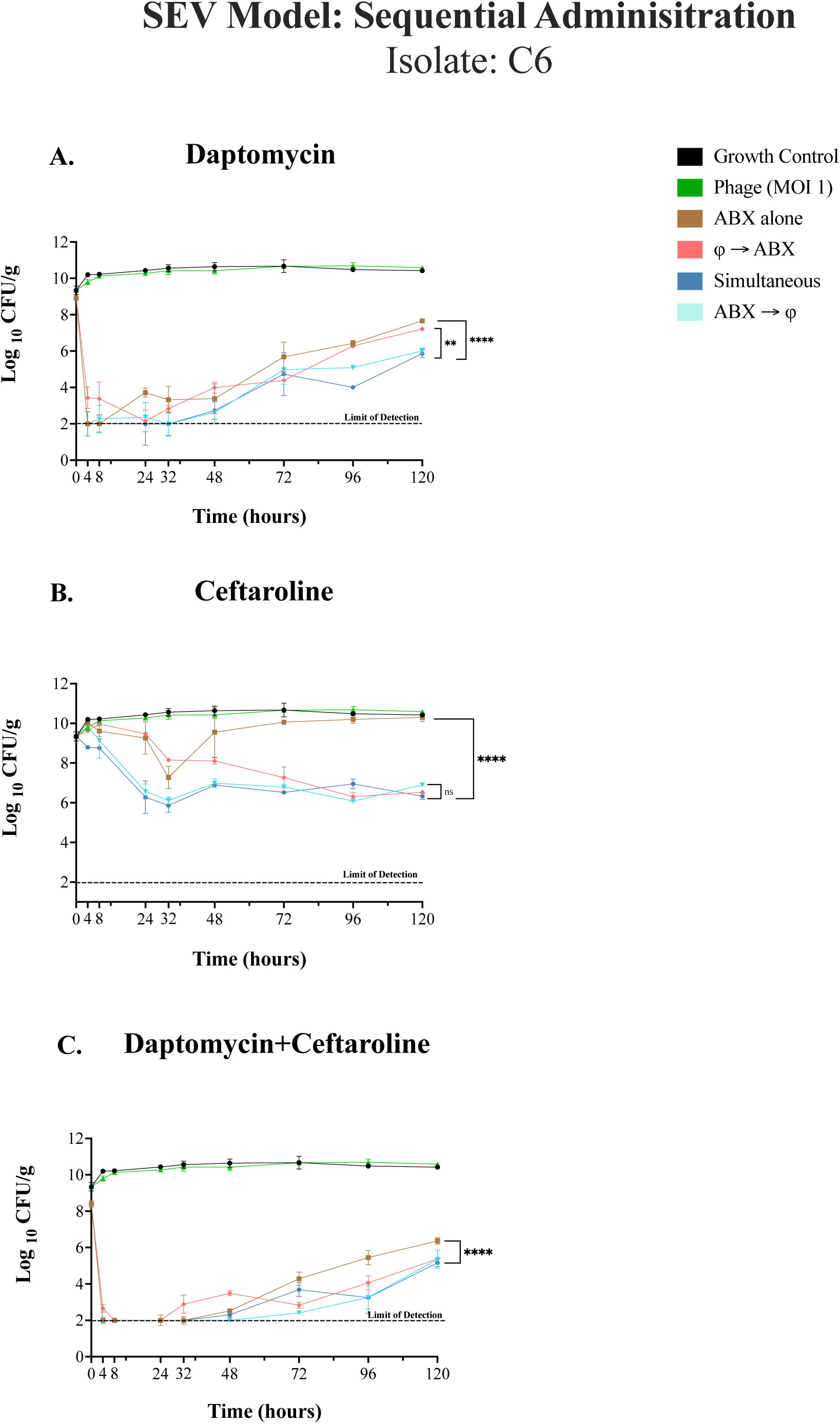
**(A-C)**. Isolate C6, *Ex-vivo* Simulated Endocardial Vegetation (SEV) Model over 120h (5 Days). The graph illustrates the change in bacterial counts (log¡ CFU/g) over a 120-hour period across different treatment regimens. A full-factorial model was used to analyze main effects (treatment regimen and MIC level) and their interaction. A two-way ANOVA was performed, followed by Tukey9s multiple comparison test for post hoc analysis, adjusting p-values to maintain statistical rigor. To account for variability across repeated measures, family-wise error rate correction was applied (³ = 0.05, 95% confidence interval). Confidence intervals were plotted for visualization, and grand means summarized trends across groups. The overall statistical significance between treatments (simultaneous vs. antibiotics alone is indicated (p = 0.0037). The dashed line indicates the limit of detection at 2.0 log_10_ CFU/g. Asterisks denote statistical significance: p < 0.05*, p < 0.01**, p < 0.001***, and p < 0.0001****. Error bars represent the standard deviation from the mean of biological replicates (quadruplicate samples/timepoint). The results provided insight into treatment efficacy across MIC levels and administration sequences, identifying statistically significant differences and the most effective regimens (p < 0.05).

Bacteriophages were added to the system every 24h. The dose (10^10^ total PFU) was chosen to achieve a target multiplicity of infection (tMOI) of 1.0 at the time of first phage addition, based on prior data regarding the average CFU concentration in the SEV and TKAs on Day 1 (Figures 2, 4-7). (17, 30) Figure 8 (A-C) and Figure 9 depict the effects of different treatment sequences involving antibiotics ceftaroline, daptomycin, and/or daptomycin + ceftaroline along with the bacteriophage cocktail on bacterial burden (log_10_ CFU/g) over 120 hours.

**Figure 9.**
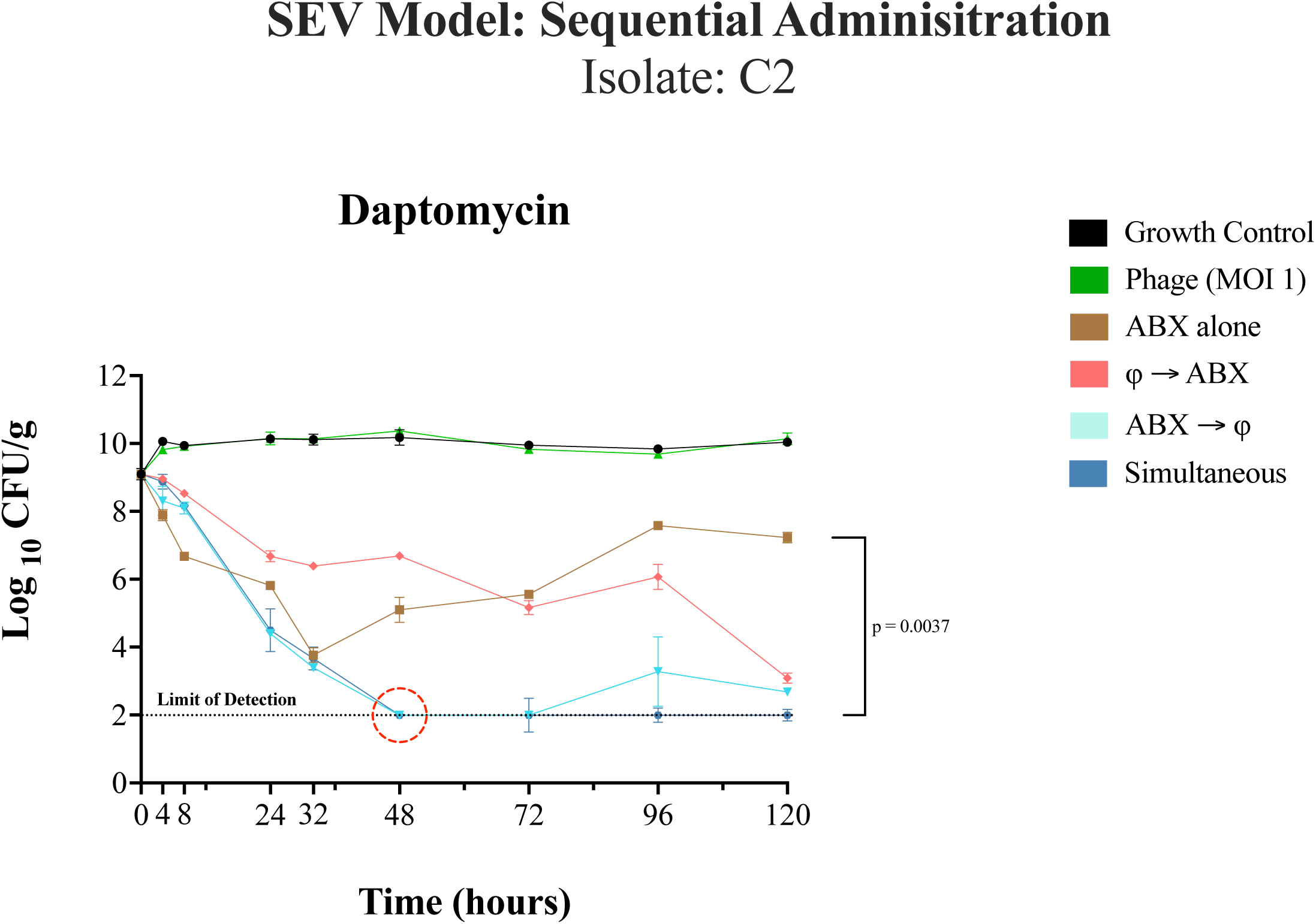
Isolate C2, *Ex-vivo* Simulated Endocardial Vegetation (SEV) Model over 120h (5 Days) using DAP-only based regimens. The graph illustrates the change in bacterial counts (log¡ CFU/g) over a 120-hour period across different treatment regimens. A full-factorial model was used to analyze main effects (treatment regimen and MIC level) and their interaction. A two-way ANOVA was performed, followed by Tukey9s multiple comparison test for post hoc analysis, adjusting p-values to maintain statistical rigor. To account for variability across repeated measures, family-wise error rate correction was applied (³ = 0.05, 95% confidence interval).Confidence intervals were plotted for visualization, and grand means summarized trends across groups.The red circle denotes time to limit of detection for simultaneous and antibiotic-first administration. The overall statistical significance between treatments (simultaneous vs. antibiotics alone is indicated (p = 0.0037). The dashed line indicates the limit of detection at 2.0 log_10_ CFU/g. Asterisks denote statistical significance: p < 0.05*, p < 0.01**, p < 0.001***, and p < 0.0001****. Error bars represent the standard deviation from the mean of biological replicates (quadruplicate samples/timepoint). The results provided insight into treatment efficacy across MIC levels and administration sequences, identifying statistically significant differences and the most effective regimens (p < 0.05).

### SEV Model: Sequential Phage and Antibiotic Administration against Isolate C6

In Figure 8A, simultaneous and daptomycin-first treatment exhibited the most pronounced bacterial reduction, significantly outperforming both the growth control and daptomycin-alone treatment regimens (p < 0.0001). Simultaneous administration, daptomycin-first, or bacteriophage-first sequences were all significantly better than the growth control but differed in efficacy, with bacteriophage-first treatment showing reduced bacterial burden compared to daptomycin-first treatment (p = 0.0029). No detectable phage-resistant mutants (<LoD) were observed in all treatment groups at the 120-hour.

Key findings at the 120-hour timepoint for Figure 8B show simultaneous treatment, bacteriophage-first, and ceftaroline-first treatment regimens significantly reduced bacterial counts compared to the growth control and antibiotics alone (p < 0.0001). The highest numerical reduction was observed with simultaneous treatment (−4.09 ±0.16 log_10_ CFU/g); however, this was not significantly different from either the antibiotic-first or phage-first treatments (p = 0.596). Treatment with either bacteriophage or ceftaroline alone, did not differ significantly from the growth control, indicating limited efficacy when administered individually. While phage-first treatment appeared slower than antibiotic-first or simultaneous administration, ceftaroline initially reduced bacterial counts but was followed by regrowth. Despite, the CPT MIC remaining the same, antibiotic-first, phage-first, and simultaneous treatments showed high phage resistance emergence (FoR = 9.12E-01, 9.83E-01, and 9.88E-01, respectively).

In Figure 8C, the combination of DAP+CPT (simultaneous, bacteriophage-first, and antibiotic-first) demonstrated significant bacterial reductions compared to the growth control and single-agent treatments (p < 0.0001). Simultaneous administration resulted in the highest numerical bacterial reduction (−5.25±0.21 log_10_ CFU/g), though this was not significantly different from antibiotic- or phage-first sequences. No significant differences were observed between bacteriophage-first and antibiotic-first treatment regimens, indicating equivalent efficacy of sequential approaches in this model. Simultaneous treatment resulted in the highest frequency of resistance (FoR = 1.66E-01), compared to antibiotic-first and phage-first strategies (FoR = 1.28E-01 and 1.21E-01, respectively). Additionally, no changes in daptomycin or ceftaroline MICs were observed across all Figures 8A-C, at the 120-hour timepoint.

### SEV Model: Sequential Phage and Antibiotic Administration against Isolate C2

In the C2 SEV model with high bacterial inoculum and humanized antibiotic concentrations (Figure 9), the untreated growth control showed stable bacterial counts around 10.0 log₁₀ CFU/g throughout the duration of the experiment, indicating robust bacterial proliferation in the absence of treatment. Using the phage cocktail alone led to an initial modest decrease in the first 24h (−1.2 log_10_ CFU/g), then plateaued around 9.0 log₁₀ CFU/g, mirroring the growth control group. Additionally, phage resistance was observed at the 12-hour timepoint (FoR = 7.62E-01).

Daptomycin alone had a pronounced reduction in bacterial counts within the first 24h (−5.0 log₁₀ CFU/g), however, bacterial counts began to increase again after 48 hours, indicating a regrowth of the bacterial population with no changes in the DAP MIC observed. Both simultaneous and antibiotic-first administration strategies achieved bacterial killing to detection limits at 48h, with simultaneous treatment maintaining bacterial counts below detection limits out to 120h. Simultaneous administration demonstrated significantly greater efficacy than daptomycin alone (p = 0.0037). No detectable phage-resistant mutants (<LoD) were observed in all sequential treatment groups at the 120-hour.

## MATERIALS AND METHODS

### Bacteriophage Selection

This study was centered on using a two-phage cocktail. Two *S. aureus* myophages with lytic properties, both closely related to phage K and sharing >95% intergenomic similarity (31) (31–33) were selected for this study based on prior phage screening experiments and methodology previously completed in our laboratory. (25, 30) The phages selected, Intesti13 and Sb-1, exhibited activity against our targeted DNS-MRSA strains in phage plaque assays. (34) Additionally, prior research demonstrated that these phages had unique host ranges, and resistance to one phage did not necessarily result in cross-resistance within the phage cocktail. (17) (26, 30) Based on these criteria, phages Sb-1 and Intesti13 were selected for in vitro, and ex vivo experiments outlined below against selected isolates. Both bacteriophages, Sb-1 (35) and Intesti13, were obtained from bacteriophage stocks procured from the Georgia Eliava Institute (Tbilisi, Georgia). In line with previously described methods, phage production and quantification of Sb-1 and Intesti13 were carried out using host strains D712 and ATCC 19685, respectively. (36)

### Bacterial Isolate Selection

This study examined a total of sixteen clinical DNS-MRSA strains isolated from bacteremic patients. Thirteen were obtained from the Cubist Pharmaceuticals Isolate Collection, Cubist Biorepository (Cambridge, MA) (37) while the remaining three isolates were provided to us: 684 (VA Hospital, Detroit, MI, USA) (38, 39) JKD6005 (Brisbane, AUS) (40, 41) and J03 (Detroit, MI, USA) (42). These strains are archived in the Rybak Anti-Infective Research Laboratory, where they have been periodically tested to confirm their viability and retention of DNS phenotypes through continuous MIC testing.

### Antimicrobial Selection and Susceptibility Testing

Each selected isolate’s minimum inhibitory concentration (MIC) against daptomycin (DAP), ceftaroline (CPT), and vancomycin (VAN), were manually tested in duplicate and validated via two methods. (i) microbroth dilution (MBD) performed at approximately 10^6^ CFU/mL according to the CLSI guidelines (ii) ETEST® (bioMérieux, FR) and Liofilm® MIC Test Strip (FisherScientific) (43) performed on Tryptic Soy Agar (TSA) (BD, Chicago, IL, USA), incubated at 37°C and read 24h later. MICs disseminated via MBD were conducted in 96-well microplates (Thermo Fisher Scientific) using Mueller-Hinton Broth (MHB; Difco, Detroit, MI) supplemented with an appropriate target of calcium [50mg/L] and magnesium [12.5mg/L] concentration (MHB50) to account for any MIC shifts when using daptomycin. (43, 44) Daptomycin (DAP) was obtained from Xellia Pharmaceuticals (Buffalo Grove, IL, USA) and ceftaroline (CPT) analytical powder was sourced from AbbVie, Inc. Chicago, IL, USA).

### Continuous Growth Suppression Screening Curves

Continuous growth suppression curves were conducted against sixteen DNS-MRSA to identify isolates that were the least susceptible to the combination of DAP+CPT during planktonic growth and therefore, candidates for combining the phage cocktail in subsequent experiments. Overnight cultures of each isolate were diluted to an initial concentration of approximately 10^6^ CFU/mL. Antibiotics were introduced into designated wells at concentrations 0.5x the MIC, based on previous susceptibility testing (Table 1). The growth suppression curves and MICs were performed in triplicate and duplicate to ensure both accuracy and reproducibility. The 24h growth assay was performed over an extended time (18-24h) using a continuous reading spectrophotometer (LogPhase 600, Agilent BioTek; Winooski, VT, USA) to monitor bacterial growth. Optical density at 600 nm (OD_600_) was measured approximately every 10 minutes to capture a comprehensive understanding of bacterial growth dynamics in response to the antibiotic treatment.

**Table 1.**
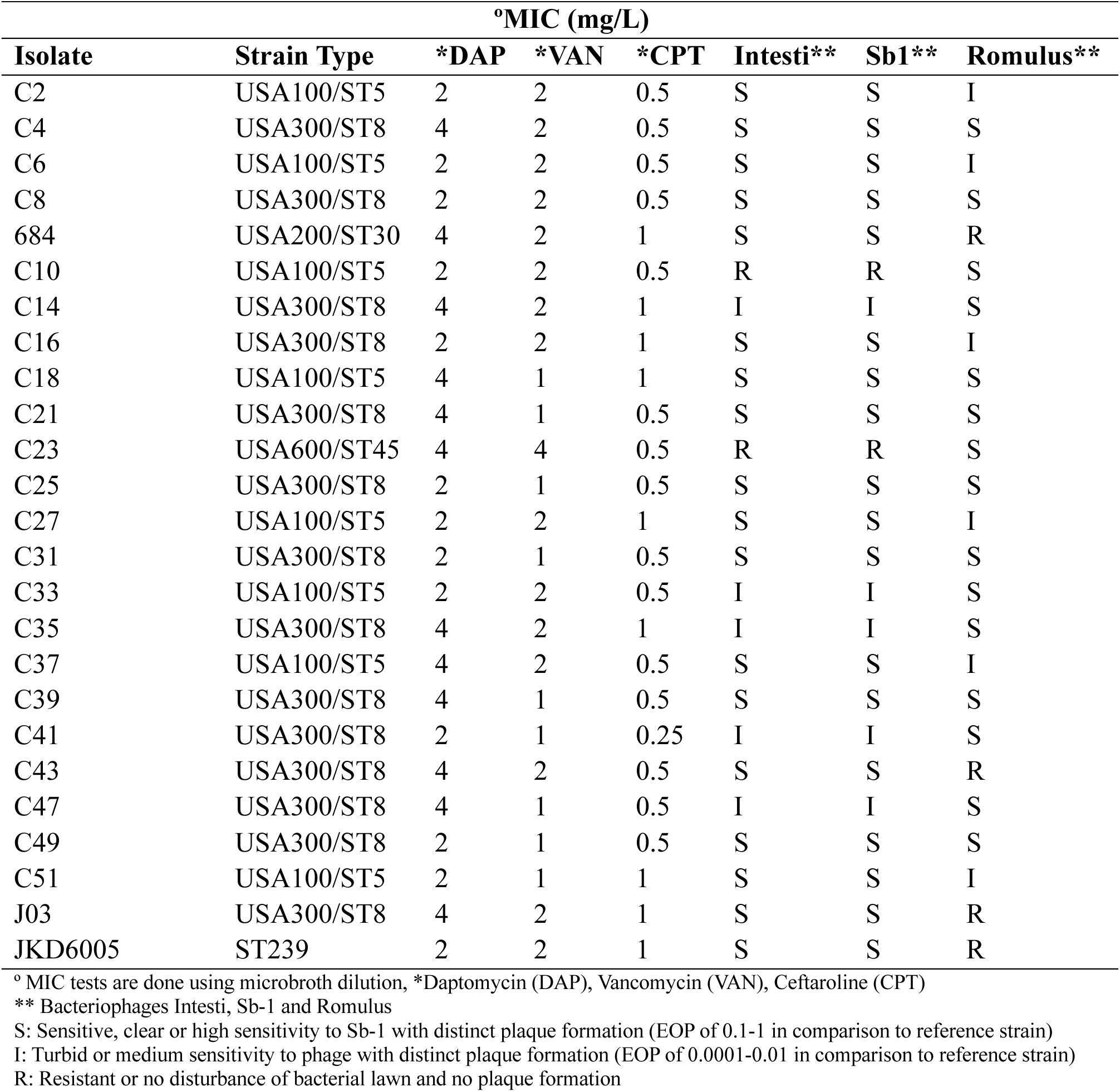
List of DNS-MRSA strains with MIC values for daptomycin (DAP), vancomycin (VAN), and ceftaroline (CPT), along with bacteriophage susceptibilities.

**Table 2.**
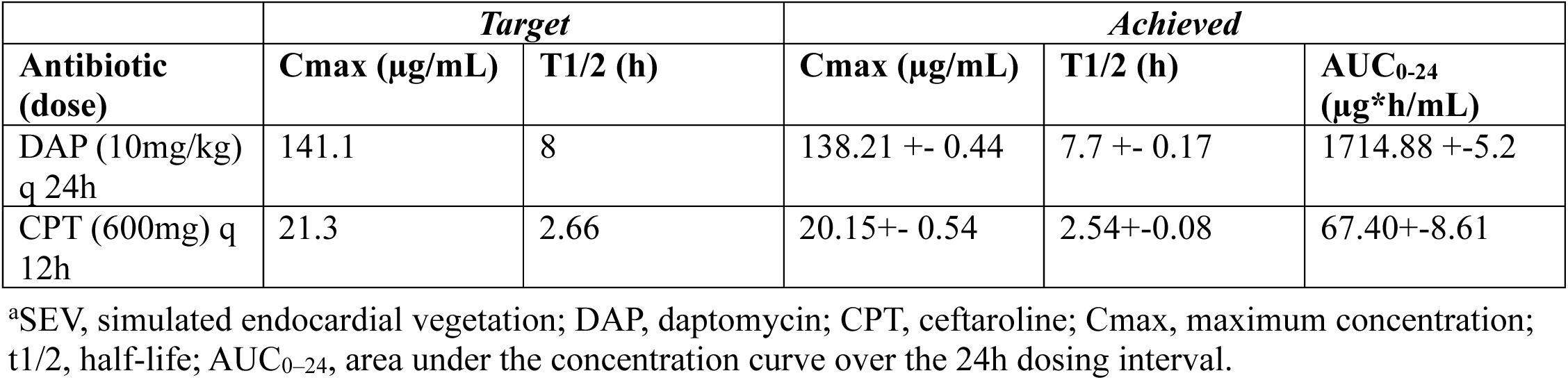
Pharmacokinetic parameters for antibiotics used in ex vivo SEV models^a^.

| Antibiotic (dose) | Target |  | Achieved |  |  |
| --- | --- | --- | --- | --- | --- |
|  | Cmax (µg/mL) | T1/2 (h) | Cmax (µg/mL) | T1/2 (h) | AUC<sub>0-24</sub> (µg*h/mL) |
| DAP (10mg/kg) q 24h | 141.1 | 8 | 138.21 +- 0.44 | 7.7 +- 0.17 | 1714.88 +-5.2 |
| CPT (600mg) q 12h | 21.3 | 2.66 | 20.15+- 0.54 | 2.54+-0.08 | 67.40+-8.61 |

<sup>a</sup>SEV, simulated endocardial vegetation; DAP, daptomycin; CPT, ceftaroline; Cmax, maximum concentration; t1/2, half-life; AUC<sub>0-24</sub>, area under the concentration curve over the 24h dosing interval.

In this study, the threshold for growth suppression was determined based on the lower bounds of the 95% confidence intervals across all strains, yielding a value of OD_600_ = 0.266. This threshold represents a conservative measure of significant suppression, accounting for variability and uncertainty in the optical density (OD) measurements. Strains for which the mean OD values remained below this threshold until at least 22h, were considered effectively suppressed under the tested conditions.

### Evaluation of Sequential Administration in Experimental Settings (in vivo and ex-vivo)

To assess the impact of the order of administration on phage-antibiotic interactions, three main sequential administration sequences were tested at the two-hour mark: (i) phage first, (ii) antibiotics first, followed by (iii) simultaneous administration. The two-hour interval was chosen based on one-step growth curve data, which suggests that it takes about two hours for the specific *Staphylococcal* phages (Intesti13 and Sb-1) to complete their lytic cycle. (45, 46) Therefore, there was hypothetically time for one phage replication cycle to occur before antibiotic addition during phage-first sequences. Sequential administration at varying MICs (0.5x, 1x, and 2x) were initially evaluated in 24h time-kill assays. Based on these results, two isolates (one USA/100 and one USA/300 strain) were selected for further testing in a 5-day, *ex-vivo* simulated endocardial vegetation (SEV) model.

### Time-kill Assays

In vitro high-inoculum (∼9.0 log_10_ CFU) 24h time-kill assays using antibiotics alone at varying MICs were conducted in 24-well microplates (Thermo Fisher Scientific; Detroit, MI) as previously described. (17) Antibiotics DAP and CPT were introduced to their designated wells in duplicate at 0.5x, 1x and 2x the MIC. Five assays were conducted in duplicate against the five DNS-MRSA isolates selected [684, C2, C6, C49, C31], chosen by growth suppression curves.

The two-phage cocktail with a multiplicity of infection (MOI) of 1 was used, as supported by prior work. (17, 25) Each well was supplemented with MHB50 and inoculated at ∼9.0 Log _10_ CFU/mL, to simulate a deep seated infection. The regimen consisting of DAP ± CPT ± the two-phage cocktail, was then applied in the remaining 24h TKAs at varying increments of the MICs to better understand clinical treatment failure concentrations and to assess whether the concentration of antibiotics at various increments of the MIC levels affect the optimal order of phage administration. The impact of order of administration was evaluated using 24h TKAs with phage ± DAP and or/CPT at 0.5x, 1x and 2x the MIC as follows: Phage first administration: Phage administered followed by antibiotic(s) two hours later., Antibiotic(s) administered first:

Antibiotic(s) administered followed by phage two hours later and Simultaneous administration: Phage and antibiotic(s) administered together.

All TKAs were incubated at 37°C for 24h, maintaining a constant shaking speed at 50RPM. 100-µL samples were ascetically removed from each well at 24h. Subsequent steps involved eliminating antibiotic and phage carryover through two rounds of centrifugation, supernatant removal, and appropriate dilutions in 0.9% saline. Samples were then plated on TSA and incubated at 37°C for 24h before bacterial colony counting (Scan 1200, Interscience for Microbiology, Saint Nom la Breteche, France; detection limit of 2.0 Log_10_ CFU/mL). Synergy was defined as a ≥2 log_10_ CFU/mL reduction compared to the most effective single regimen. Bactericidal activity was defined as a ≥3 log_10_ CFU/mL reduction from most effective single regimen.

### Ex-vivo Pharmacokinetic/Pharmacodynamic (PK/PD) Simulated Endocardial Vegetation (SEV) Model

*Ex-vivo* SEV PK/PD models were conducted in duplicate, with SEV clots prepared following established methods (17, 38, 47) The models were maintained at 37°C for the full 120-hour duration. To simulate antibiotic half-lives, MHB50 was cycled in and out at a controlled rate, with humanized doses of DAP (10 mg/kg) administered every 24 hours. Phages were added to the SEV model every 24h. The dose (10^10^ total PFU) was chosen to achieve a target multiplicity of infection (tMOI) of 1.0 at the time of first phage addition. SEV samples were collected in duplicate at designated time points (0, 4, 8, 24, 32, 48, 72, 96, and 120 hours), totaling four samples per model. Each SEV clot underwent homogenization, followed by two rounds of centrifugation. After each centrifugation step, the supernatant was removed and replaced with normal saline to eliminate residual antibiotic and phage carryover, following established protocols. (17)

### Pharmacokinetic Analysis

Antibiotic pharmacokinetic samples were collected in duplicate from the injection port of each SEV infection model at specified time points (0, 4, 8, 24, 32, 48, 72, 96, and 120 hours) to confirm target antibiotic concentrations. As previously outlined, all collected samples were stored at −80°C until analysis (17, 38, 48) DAP concentrations were determined using a validated HPLC assay, meeting College of American Pathologists’ standards, with an intraday coefficient of variance below 2% across all concentration levels. Pharmacokinetic parameters, including half-life, peak concentration, and area under the curve (AUC) (calculated using the trapezoidal method), were determined using PK Analyst software (version 1.10; MicroMath Scientific Software, Salt Lake City, UT, USA) (17)

CPT concentrations were analyzed through bioassay, utilizing *Bacillus subtilis* ATCC 6633 as the test organism. For this, blank 1/4-inch disks were impregnated with 10 µL of either standard solutions or test samples. Standards were tested in duplicate by placing disks on agar plates (antibiotic medium 11) pre-inoculated with a 0.5 McFarland suspension of the bacterial strain. The bioassay exhibited an intraday coefficient of variance below 4.7% for high, medium, and low broth standards. Plates were incubated at 37°C for 24 hours, after which inhibition zones were measured using a ProtoCOL plate reader (Microbiology International, Frederick, MD, USA) (17) A complete table outlining the target and achieved pharmacokinetic parameters for DAP± CPT used in the *ex-vivo* models can be found in Table 2.

### Post-SEV Antibiotic and Phage Susceptibility Testing

#### Antibiotic susceptibility

Changes in antibiotic susceptibility compared to baseline were evaluated in each final SEV timepoint sample (120h) using specific antibiotic-embedded agar via MBD. 100μL of the 120h SEV sample was plated onto individual TSA plates containing threefold the baseline MIC of the drug (DAP/CPT) used in the model. Plates were examined for growth after 24h and 48h of incubation at 36°C. MIC changes via MBD according to CLSI guidelines were performed if growth was present. (43) For either assay, if samples demonstrated MIC changes of ≥2 dilution from baseline (elevation or reduction in MIC), then they were passed for a 3-day consecutive period with MIC testing each day. For samples maintaining the ≥2 dilution changes in MIC from baseline following the 3-day pass, additional SEV samples were assessed for resistance in a backward stepwise manner from 96h to earlier time points until a ≤1 dilution change in MIC was identified for the sample. (17)

#### Phage susceptibility

Bacterial samples from phage-treated conditions, were thawed for susceptibility testing. For each sample, 10µL of bacterial suspension was combined with 100 µL of high-titer phage stock (≥10^10^ PFU/mL) in a labeled snap-cap tube. The mixture was incubated in a shaker incubator for 10 minutes to allow for phage adsorption. Meanwhile, 0.7% HIB agar overlay was prepared, and the temperature was measured to ensure it remained within the 45– 50°C. (36) After incubation, 3mL of HIB agar overlay was added to each bacterial-phage mixture and poured onto pre-prepared agar plates (1.5% HIB). Plates were incubated at 36°C overnight. Following 24 hours of incubation, plates were visually inspected for colonies (bacteriophage-insensitive mutants, aka BIMs). BIMs were counted using a Biotek plate reader, and results were recorded. Plates were left at room temperature for an additional 24 hours. After a total of 48 hours (24 hours at 36°C + 24 hours at room temperature), BIM counts were re-evaluated using the plate reader (t48 BIMs), and data recorded. If necessary, plates were stored at 4°C until further validation via the double drop method similar to that previously described. (49) The apparent frequency of resistance (FoR) was calculated by the total CFU/mL used to create the resistance plate divided by total average number of bacteriophage insensitive mutants (BMIs) at 48h. (17, 34)

### Statistical Analysis

Statistical analyses were performed on data generated from experiments conducted in at least two replicates and repeated twice, with results presented as mean ± standard deviation (n = 4), unless otherwise indicated. Analyses were conducted using SPSS version 29 (IBM Corp., Armonk, NY, USA) and GraphPad Prism software (versions 9.2.0 and 10.0; GraphPad, La Jolla, CA, USA). Differences in bacterial burden reduction between phage and/or antibiotic regimens were assessed using one-way ANOVA followed by Tukey’s multiple comparisons test, with statistical significance set at p < 0.05.

For the 24h continuous growth suppression data, a two-way repeated measures ANOVA analysis with Tukey’s HSD post hoc test for multiple comparisons (p<0.005) was performed to evaluate differences in bacterial growth suppression via spectrophotometric analysis (Δ OD_600_). The analysis accounted for repeated measures across time and included corrections for violations of sphericity using the Geisser-Greenhouse correction. Post hoc pairwise comparisons (Tukey’s HSD test) were conducted to further explore significant interactions at individual timepoints (S1, S2).

In 24h TKAs, to evaluate the effects of treatment regimens on bacterial reductions across different conditions, a two-way analysis of variance (ANOVA) with repeated measures was performed. The analysis considered two factors: administration sequence (i.e., phage-first, antibiotic-first, simultaneous) and minimum inhibitory concentration (MIC) levels (e.g., 0.5x, 1x, and 2x MIC). The data were grouped by administration sequence (columns) and MIC levels (rows) to assess both main effects and interaction effects between the two factors.

A full-factorial model was employed to analyze column effects, row effects, and their interaction. Tukey’s multiple comparison test was used for post hoc analysis to identify significant differences between treatment groups while controlling for multiple comparisons. Adjusted p-values were reported to ensure statistical rigor. To account for variability across repeated measures and to ensure accurate comparisons, the analysis applied family-wise error rate correction with a 95% confidence interval (alpha = 0.05). Confidence intervals were graphed for visualization, and grand means were displayed to summarize trends across groups.

The results from the two-way ANOVA, combined with pairwise comparisons, provided insight into the efficacy of treatment strategies across MIC levels and administration sequences, identifying statistically significant differences and the most effective treatment regimens (*p* < 0.05). (17)

## DISCUSSION

### Phage-Antibiotic Synergy and Optimization of Administration Order

Our strain collection incorporates variability in the susceptibility of USA100/ST5 and USA300/ST8 strain types to the combination of daptomycin (DAP) and ceftaroline (CPT). In all tested experimental settings, DAP+CPT was not consistently effective across all strains. As in previous studies, however, the triple combination of DAP+CPT+phage largely overcame this strain variability and showed considerable potential in reducing bacterial counts within planktonic *Staphylococcus aureus* populations, particularly at higher antibiotic concentrations. (50–54) Given the range of results that have been reported in order of administration studies with staphylococcal phages and antibiotics, we’ve endeavored to look for trends that carry through from simple planktonic growth suppression experiments through to more complex *ex-vivo* models with humanized antibiotic PK. In general, our results confirm and expand on our previous findings that simultaneous bacteriophage-antibiotic treatments can act synergistically to improve both the magnitude and consistency of bactericidal outcomes in multiple model systems. For the most part, sequential treatments yielded comparable outcomes, however a few deviations from this trend are noteworthy.

When the order of administration made a difference in TKAs, it was typically the “antibiotic-first” regimen that tended to underperform, although this could be overcome by higher antibiotic concentrations. One hypothesis for this observation would be that administering antibiotics first could limit the availability of bacterial hosts for phage infection to occur, thus decreasing the synergistic impact of PACs. Related to this, antagonism has sometimes been shown to occur with specific antibiotics, such as protein synthesis inhibitors, presumably because phages rely on bacterial cellular machinery for replication (12, 55–57) Under these hypotheses, we would expect i) treatment order differences to disappear as the antibiotic concentration increased (which we did observe), and ii) phage-antibiotic synergy to disappear if the antibiotic concentration became sufficiently high (which we did not observe within the range of concentrations that we tested). These observations highlight ongoing questions about whether sequential administration is more effective depending on the specific antibiotic’s mechanism of action and whether the distinction between concentration-dependent and time-dependent antibiotics plays a role.

In C6, the SEV models showed no clear advantage of simultaneous over antibiotic-first administration, as both approaches performed similarly. However, one of the most dramatic instances in which the order of administration affected outcomes was in SEV models with isolate C2, where daptomycin followed by phage completely or almost completely suppressed bacterial growth within 48 h, but administering phage first before daptomycin did not approach the limit of detection until approximately 120 h (Figure 9).

Our data highlight the importance of continuing to investigate when and how the timing and sequence of bacteriophage-antibiotic combinations might affect therapeutic outcomes, particularly in infections like infective endocarditis, where bacterial persistence and resistance pose significant clinical challenges.

### Emergence of Phage Resistance in the SEV Model

A key observation in the *ex-vivo* SEV model was the emergence of phage resistance at hour-120, particularly seen in isolate C6, when treated with CPT or DAP+CPT. In contrast, no phage resistance was observed when using daptomycin alone (Table 5). Despite not reaching the limit of detection, no significant changes in daptomycin or ceftaroline MICs were observed in all treatment groups (Table 3-4). The combination of DAP+CPT+Phage (Figure 8C) performed similarly to DAP+Phage (Figure 8A), even though DAP+CPT alone demonstrated improved bacterial clearance in the absence of phage. This suggests that the addition of CPT may have facilitated the emergence of bacteriophage-insensitive mutants (BIMs), which could have limited the overall effectiveness of the combination therapy. The selective pressure exerted by CPT, rather than a saturation of bactericidal activity, likely played a role in resistance development, highlighting a key challenge in phage-antibiotic combination strategies.

**Table 3.** Antibiotic MIC for DNS-MRSA isolate C6 at baseline and in T120h SEV samples^a^.

| **Bacterial Sample** | **Antibiotic MIC (µg/mL)** |  |
| --- | --- | --- |
|  | **DAP\*** | **CPT\*** |
| Baseline (Pre-antibiotic/phage exposure) | **2-4** | **0.05** |
| End of Study: CPT |  |  |
| Growth Control | **2** | **0.05** |
| (ABX alone) CPT 600mg |  | **0.05** |
| (ABX first) CPT 600mg → Phage |  | **0.05** |
| (Phage first) CPT Phage → CPT 600mg |  | **0.05** |
| (Simultaneous) CPT 600mg + Phage |  | **0.05** |
| End of Study: DAP |  |  |
| Growth Control | **2** |  |
| (ABX alone) DAP 10mg/kg | **2-4** |  |
| (ABX first) DAP 10mg/kg → Phage | **2-4** |  |
| (Phage first) Phage → DAP 10mg/kg | **2-4** |  |
| (Simultaneous) DAP 10mg/kg + Phage | **2-4** |  |
| End of Study: DAP+CPT |  |  |
| Growth Control | **2** | **0.05** |
| (ABX alone) DAP 10mg/kg + CPT 600mg | **2** | **0.05** |
| (ABX first) DAP 10mg/kg +CPT 600 mg → Phage | **2** | **0.05** |
| (Phage first) Phage → DAP 10mg/kg +CPT 600 mg | **2** | **0.05** |
| (Simultaneous) DAP 10mg/kg +CPT 600 mg + Phage | **2** | **0.05** |

<sup>a</sup>SEV, simulated endocardial vegetation; DAP, daptomycin; CPT, ceftaroline
\*All MICs in the table are expressed as µg/mL

**Table 4.**
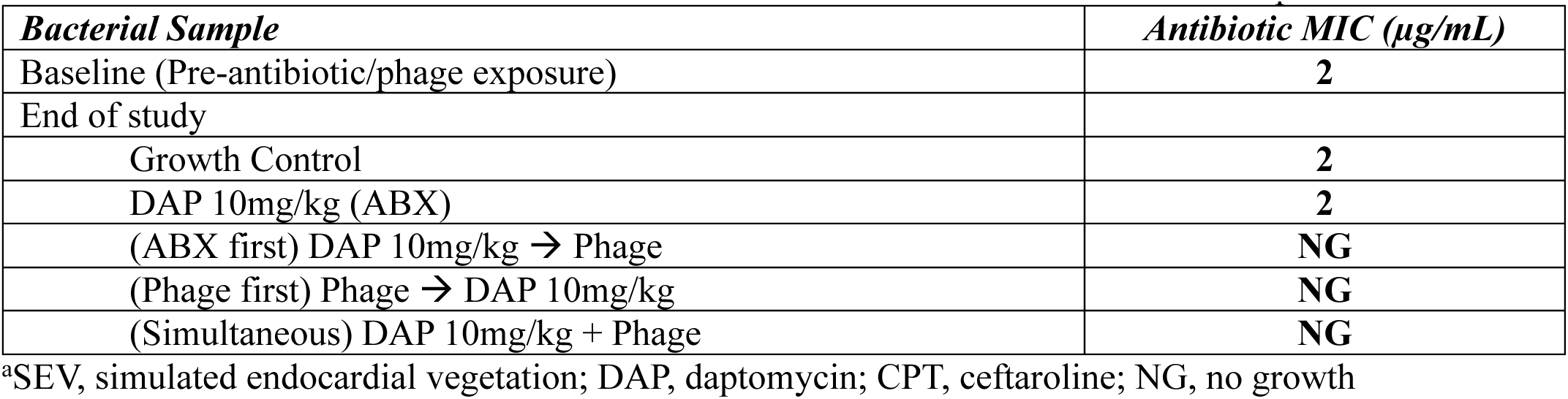
Antibiotic MIC for DNS-MRSA isolate C2 at baseline and in T120h SEV samples^a^.

| **Bacterial Sample** | **Antibiotic MIC (µg/mL)** |
| --- | --- |
| Baseline (Pre-antibiotic/phage exposure) | **2** |
| End of study |  |
| Growth Control | **2** |
| DAP 10mg/kg (ABX) | **2** |
| (ABX first) DAP 10mg/kg → Phage | **NG** |
| (Phage first) Phage → DAP 10mg/kg | **NG** |
| (Simultaneous) DAP 10mg/kg + Phage | **NG** |

<sup>a</sup>SEV, simulated endocardial vegetation; DAP, daptomycin; CPT, ceftaroline; NG, no growth

**Table 5.** Apparent frequency of resistance in DNS-MRSA strains C6 and C2 recovered from SEV models at the 120-h timepoint^,a,b,c^.

| Strain | Treatment | CFU/mL at 120-h SEV timepoint | Average Bacteriophage Insensitive Mutants (BIMS)<sup>d</sup> | Apparent FoR |
| --- | --- | --- | --- | --- |
| C6 | **DAPTOMYCIN (Fig 8A)** |  |  |  |
|  | (ABX first) DAP 10mg/kg → Phage | 4.49E+02 | 5.00E-01 | <LoD |
|  | (Phage first) Phage → DAP 10mg/kg | 5.26E+02 | 2.50E+00 | <LoD |
|  | (Simultaneous) DAP 10mg/kg + Phage | 3.59E+02 | 0 | <LoD |
|  | **CEFTAROLINE (Fig 8B)** |  |  |  |
|  | (ABX first) CPT 600mg → Phage | 4.55E+02 | 4.15E+02 | 9.12E-01 |
|  | (Phage first) Phage → CPT 600mg | 3.91E+02 | 3.84E+02 | 9.83E-01 |
|  | (Simultaneous) CPT 600mg + Phage | 3.86E+02 | 3.81E+02 | 9.88E-01 |
|  | **DAPTOMYCIN+CEFTAROLINE (Fig 8C)** |  |  |  |
|  | (ABX first) DAP 10mg/kg + CPT 600mg → Phage | 3.42E+02 | 4.40E+01 | 1.28E-01 |
|  | (Phage first) Phage → DAP 10mg/kg + CPT 600mg | 3.43E+02 | 4.15E+01 | 1.21E-01 |
|  | (Simultaneous) DAP 10mg/kg + CPT 600mg + Phage | 3.08E+02 | 5.10E+01 | 1.66E-01 |
|  | **PHAGE ALONE (Fig 8 A-C)** |  |  |  |
|  | Intesti13+Sb-1 | 9.62E+02 | 9.50E+02 | 9.88E-01 |

| Strain | Treatment | CFU/mL | Average Bacteriophage Insensitive Mutants (BIMS) at 48h | Apparent FoR |
| --- | --- | --- | --- | --- |
| **C2** | **DAPTOMYCIN (Fig 8A)** |  |  |  |
|  | (ABX first) DAP 10mg/kg → Phage | 2.70E+02 | 0 | <LoD |
|  | (Phage first) Phage → DAP 10mg/kg | 3.29E+02 | 5.00E+00 | <LoD |
|  | (Simultaneous) DAP 10mg/kg + Phage | 2.39E+02 | 0 | <LoD |
|  | **PHAGE ALONE (Fig 8 A-C)** |  |  |  |
|  | Intesti13+Sb-1 | 9.19E+02 | 7.00E+02 | 7.62E-01 |

<sup>a</sup>Abbreviations: CFU, colony-forming unit; FoR, frequency of resistance; LoD, limit of detection; DAP, daptomycin; CPT, ceftaroline.
<sup>b</sup>FoR: [colonies that arise on the resistance plate]/[CFU per milliliter used to create the plate].
<sup>c</sup>LoD: If <LoD is indicated, then the apparent FoR equals 0.
<sup>d</sup>Counted after 48 h incubation on the resistance plate

### Role of Genetic Mutations in Phage Resistance

The CUBIST-2 (C2) (58) and CUBIST-6 (C6) (59) strains exhibited distinct resistance mechanisms influencing DAP and phage activity. In C2, the presence of the *mprF* L826F mutation is associated with daptomycin resistance by altering membrane charge, thereby reducing DAP’s effectiveness. (60, 61) However, current research has shown that this mutation does not directly impact phage receptors, allowing phage sensitivity to remain intact when DAP is combined with phage alone. (15) In contrast, C6 harbors additional mutations in *walK* (L494V) and *cls2* (F60S) alongside *mprF* (T345A), which may further complicate antibiotic-phage interactions. Mutations in *walK* impact cell wall stress responses, potentially modifying surface structures like wall teichoic acids (WTAs) that serve as phage attachment sites. (62, 63) These mutations may exacerbate resistance development when multiple stressors (phage±DAP/CPT) are applied, as bacteria adapt their surface receptors under selective pressure.

Beyond genetic mutations, CPT itself may drive collateral phage resistance through its effects on WTAs and lipoteichoic acids (LTAs). By targeting penicillin-binding proteins (PBPs), particularly PBP2a, CPT disrupts peptidoglycan synthesis, triggering compensatory mechanisms to stabilize the bacterial cell wall. These compensatory adaptations include alterations in WTA structure, such as changes in chain length or glycosylation patterns, as well as modifications to LTA to maintain membrane anchoring. (64) Since K-like staphylococcal myophages bind to the WTA backbone (65), these CPT-induced modifications might negatively impact phage binding efficiency, conferring collateral phage resistance as outlined in Table 5. (66)

This may explain why phage resistance emerged in the SEV model at T120, particularly in C6 (Figure 8B). In CPT-only conditions, bacteria may have had more time to adapt their WTA/LTA structure, thereby reducing phage susceptibility. However, in DAP+CPT regimens, the additional membrane stress from daptomycin may have overwhelmed bacterial survival pathways, limiting opportunities for compensatory WTA/LTA modifications. These findings highlight the complex interplay between antibiotic pressure, phage susceptibility, and bacterial adaptation mechanisms in DNS-MRSA strains.

### Future Work and Clinical Implications

Limitations of the study include the narrow scope of the phage cocktail used, as only two phages were tested in combination. This may limit the generalizability of the findings, particularly where the two-phage cocktail failed to reach the limit of detection (LOD) by T120 (SEV *ex-vivo* models) or T24h (24h TKAs). In the SEV *ex-vivo* model, phage resistance was observed by T120, particularly with isolate C6 (Table 5) as well as the combination of DAP+CPT+Phage not reaching the limit of detection, indicating limitations in the efficacy of the two-phage cocktail used. To address this, we explored the addition of Romulus, a myovirus previously shown to mitigate resistance in DNS-MRSA strains when combined with DAP+/CPT+/2-phage cocktail (Sb-1 and Intesti13) (17, 30), perhaps because it has better activity than Intesti13 and Sb-1 against USA300 MRSA at 37°C. (31)

In 24h TKAs, the addition of Romulus to the two-phage cocktail demonstrated enhanced bacterial killing in isolate C31 compared to C6. This finding may be attributed to C31’s susceptibility to Romulus, whereas C6 is resistant, limiting the effectiveness of Romulus when added to the two-phage cocktail. Supplementary Figures S4 and S5 present the comparative results of the three-phage cocktail (including Romulus) versus the two-phage cocktail, as well as an evaluation of sequential administration sequences in C6 (S6). Based on these findings, future work will focus on sequential administration strategies incorporating the three-phage cocktail to evaluate its potential impact on mitigating resistance and enhance bacterial reduction.

Another limitation was the lack of CFU/mL assessment at the 2-hour time point, just before administering either phage or antibiotics. Determining the bacterial inoculum at this stage could provide valuable insights into the effectiveness of initial phage or antibiotic treatment during the first two hours, before introducing an additional agent.

Our research emphasizes the need for strain-specific evaluations to optimize bacteriophage-antibiotic therapies, particularly when considering variability in susceptibility among USA100/ST5 and USA300/ST8 isolates. Linking these susceptibility patterns to genetic characteristics could further aid in refining treatment strategies and mitigating resistance in challenging infections. Moreover, CPT-induced WTA/LTA alterations may lead to collateral phage resistance, particularly when bacterial cells compensate for cell wall stress. Understanding these molecular interactions between phage receptors and antibiotic-induced structural adaptations will be crucial in designing more effective phage-antibiotic combination therapies for DNS-MRSA infections. Expanding these investigations has the potential to refine treatment strategies, enhance efficacy, and minimize resistance development, while also clarifying the clinical relevance of early phage-antibiotic administration in real-world scenarios.

## CONCLUSION

In conclusion, this study provides critical insights into the potential of bacteriophage-antibiotic combinations to combat resistant *Staphylococcus aureus* infections, emphasizing the importance of both the timing and sequence of administration. The findings demonstrate the enhanced activity of simultaneous phage-antibiotic treatments and highlight the influence of sequential strategies, particularly when considering concentration-dependent and time-dependent antibiotics. Variability in phage susceptibility across isolates underscores the necessity of strain-specific evaluations and the integration of genomic insights to optimize therapeutic strategies when using PACs. While the addition of Romulus to the phage cocktail shows promise in mitigating resistance under specific conditions, limitations in phage efficacy and resistance emergence point to the need for continued refinement of these approaches. Our future research will focus on expanding the scope of phage cocktails and exploring sequential administration strategies to better understand the potential of phage-antibiotic therapies.

Author order was determined based on the magnitude of each author’s contribution to the research. The first author made the most significant contributions, and Subsequent authors are listed in descending order of their involvement in the study. The last author served as the senior supervisor and principal investigator, providing guidance and oversight throughout the research process.

**C.B.**: conceptualization, methodology, data curation, writing (original and final draft), visualization, supervision, and project administration

**S.V.:** data curation, formal analysis, writing (review and editing)

**R.R.**: data curation, writing (review and editing)

**A. B**.: bioinformatics, writing (review and editing)

**S.L.:** investigation, methodology, conceptualization, writing (review and editing)

**A.B.**: Review and editing original draft, resources

**M.R.:** principal investigator; conceptualization, methodology, writing (original and final draft), visualization, supervision, project administration and funding support.

